# Pharmacokinetics Of Transbuccal Swab-Administered Naloxone-HCl Using A Novel Plant-Based Resin Formulation In Dogs

**DOI:** 10.1101/2020.12.07.415372

**Authors:** Randolph M. Johnson, Nooshin T. Azimi, Edward F. Schnipper

**Affiliations:** Statim Pharmaceuticals, Inc. Menlo Park, CA, U.S.A.; Angarus Therapeutics, Inc. Portola Valley, CA, U.S.A.

## Abstract

A proof-of-concept transbuccal swab delivery of naloxone-HCL study using a mucoadhesive, plant-based film-forming resin formulation demonstrated comparable blood levels to benchmark intramuscular (IM) injection in highly predictive dog model. Results from this study allow the potential to translate rapid onset in humans with therapeutic blood levels being reached in 2-3 minutes comparable to that observed with commercially available parenteral injections and intranasal administrations. The simplicity, ease of delivery and rapid effectiveness has the potential to meet the public health emergency needs in the rescue of opioid overdosing.

## Introduction

Naloxone is a potent μ-opioid receptor competitive antagonist and the benchmarked FDA-approved standard-of-care antidote to opioid overdose. Currently naloxone is administered *via* the intramuscular (IM), intravenous (IV), subcutaneous (SC) and intranasal (IN) routes of administration.^1–5^ All routes deliver naloxone in an effective and life-saving manner yet there remain drawbacks within use by first-responders, at-home family members and public laypersons.^6,7^

Due to the ease of administration, needle-free routes of delivery are attractive to patients and health care providers. Herein, we describe the formulation and pharmacokinetics of naloxone for simple, needle-free delivery *via* the buccal cavity and compare the pharmacokinetics of the new formulations with an IM administration of naloxone.

The buccal route of drug delivery has many advantages for systemic drug absorption.^8–10^ The noninvasive nature of administration, the ease, precision and convenience of dosing, the increased permeability of the non-keratinized buccal mucosa, and the mucosa’s dense vasculature make this a suitable route of drug delivery for therapeutic indications needing rapid systemic absorption of life-saving drugs in life-threatening emergencies.^11,12^ Drugs absorbed through the buccal mucosa enter the body *via* the jugular vein, bypassing first-pass liver metabolism and gastric/intestinal enzyme–mediated degradation.^10^ The challenges of buccal route of delivery are primarily related to overcoming the 70 micron mucus-coating barrier and the constant daily production and turnover of 750-1000 mL of saliva which causes a high clearance of drugs from the oral cavity.^13–17^

Botanical-sourced benzoin gum resins as tinctures are used commercially over-the-counter in the medical arts for enhancing adherence of surgical bandages as well as mucosal adhesive properties for use as oral mucosal protectant applications.^18–21^ We sought to test the feasibility of tincture of Styrax benzoin (TOB) resin, exploiting its mucoadhesive properties, to deliver naloxone systemically. Here we report a PK proof-of-concept transbuccal delivery of naloxone-HCl formulated with commercially available alcoholic tincture of Styrax benzoin swabbed onto the buccal mucosal surface of dogs, a standard animal for evaluating transbuccal systemic delivery of drugs in humans.^16,17^ The results demonstrate rapid systemic absorption with pharmacokinetic parameters and blood levels equivalent to benchmark intramuscular injection or other currently used routes of administration.

## Materials and Methods

### Preparation and Application of Naloxone-HCl Formulations for Buccal Administration

Benzoin (Styrax Benzoin) gum tincture was purchased from Hawaii Pharm LLC, Honolulu, HI. Naloxone-HCl was purchased from Spectrum Chemicals. Single dose vials of Naloxone for Injection (0.4 mg/mL Naloxone HCl) were purchased from Medline SKU 0409-1215-01H.

Dosing solutions of 4.0 mg/mL and 40.0 mg/mL naloxone in tincture of benzoin (TOB) solution were prepared by Vanton Research Laboratory, Concord, CA. Concentrations and stability of dosing solutions were confirmed by Integrated Analytical Labs, Berkeley, CA, using reverse phase HPLC Shimadzu VP Series 10 system and UV detection at 270 nm.

Puritan Sterile HydraFlock® Buccal Swab Applicators (25-3506-H) were used to apply naloxone to buccal surface. HydraFlock® was chosen as the applicator due to its attributes of high liquid absorption capacity coupled with complete release elution characteristics.

### Experimental Design, Method of Administration and Sampling Schedule

The pharmacokinetic study was performed in 3 adult, male, beagle dogs by an AAALAC-accredited tested facility at PMI, San Carlos, CA as per the USDA Animal Care Policies. Animals were fasted approximately 12 hours pre-procedures and allowed free access to water during that period. Physical examination together with blood analysis of complete blood count and serum chemistry on the day of the procedure were performed to ensure dogs were in good health. A predose sample was also collected prior to naloxone administration.

A three way-crossover study design was used to minimize number of animals. All animals were washed out of any other drug administrations for one month prior to study initiation. Each dog received each of the two (4.0 mg/mL and 40.0 mg/mL) naloxone buccal dosage solution and an IM injection (0.4 mg/mL) of naloxone on successive weeks with a one-week rest period between dosing. The benchmark dose for IM used by emergency responders is 0.4 mg/ml. Hence this dose was chosen for comparison. Naloxone was administered in the morning to ensure test animals were adequately monitored post transbuccal absorption.

Two hundred microliters (200 μL) of either 4.0 mg/mL (0.8 mg) or 40.0 mg/mL (8.0 mg) naloxone-HCl in TOB was transferred by pipette to a Puritan HydraFlock® swab applicator tip. Holding the polystyrene handle the swab applicator tip was then immediately applied to right inside cheek mucosa (buccal) with a side-to-side swabbing motion while rotating the handle of the applicator for 5-10 secs allowing complete evacuation of entire formulation onto the buccal mucosal surface. The used applicator tip was saved and stored at −20°C for analysis for residual naloxone after study completion. Results of analysis showed less than 5% of naloxone remained on the applicator tip demonstrating > 95% of naloxone was administered via swabbing application (data not shown). Animals receiving intramuscular injections were administered 1.0 mL of 0.4 mg Naloxone for Injections from single use vials.

### Sample Collection Schedule

An IV catheter was placed in the cephalic vein to collect PK blood samples and used to administer rescue medications and/or IV fluids in case a test animal experienced severe side effects of naloxone administration. Each group of animals had a ten-point sampling schedule which included predose, 5, 10, 15, 20, 30, 60, 120, 240- and 360-min time points.

Blood samples were collected from a leg vein and transferred into prechilled microcentrifuge tubes containing 2 μL of K_2_EDTA (0.5M) as anti-coagulant, placed on wet ice and processed for plasma by centrifugation at approximately 4°C, 3,000 g for 15 min within 30 min of collection. All plasma samples were then transferred into pre-labeled polypropylene microcentrifuge tubes, quick-frozen over dry ice and stored at −70°C until analysis by LC/MS/MS by Integrated Analytical Labs, Berkeley, CA.

### Measurement of Naloxone in Dog Plasma Samples

Briefly, samples were deproteinated using 40 mL sample plus 2 volumes of ice cold internal standard solution acetonitrile. Samples were centrifuged at 6100 × g for 30 min. Aliquots of each supernatant were transferred to the autosampler plate. Samples were analyzed using a Shimadzu VP series 10 HPLC system with a 20 × 2 mm Proto 200 C18 Higgins Analytical column with a gradient mobile phase of 0.2% formic acid in water or methanol. MS/MS was an Applied Biosystems/MDS SCIEX API 4000, with software analyst v.1.5. Ionization method was electrospray positive with source temperature TurbolonSpray (ESI) at 40°C using deuterium-labeled naloxone, naloxone-d5, as internal standard. The LLOQ was 0.5 ng/mL with accepted curve range of 0.5-1,000 ng/mL.

### Data Analysis and Reporting

Non-compartmental PK analysis and calculation of standard PK parameters for naloxone was performed by IAS using PKSolver For Excel.

## Results

### Naloxone-HCl in benzoin gum resin tincture applied to the buccal membrane demonstrates rapid systemic absorption

The time to maximum plasma concentration of naloxone applied at 0.8 mg (15 min) and 8.0 mg (18 min) *via* transbuccal swab displayed a comparable time not significantly different to that observed with an IM injection of 0.4 mg (22 min) (Figs. 1 & 2). Naloxone in TOB maintained mucoadhesive residence-time to allow rapid permeation and systemic absorption for up to 60 min (Fig 1 and inset). More sustained delivery occurred with the higher dose of 8.0 mg while the 0.8 mg dose was more equivalent to IM injection of 0.4 mg (Fig. 1). The greater amounts of transbucally applied naloxone (0.8 and 8.0 mg) compared to 0.4 mg IM, displayed higher Cmax levels (20.0 and 47.2 ng/mL, respectively) versus 8.1 ng/mL with 0.4 mg IM (Fig. 3). Additionally, these values are markedly higher than 9.3 ng/mL reported using a 4 mg IN fixed-dose naloxone atomizer in dogs (18). The systemic exposure represented by AUC of 8.0 mg dose is 5.3-fold higher than 0.4 mg IM in dogs, 4.3-fold higher than reported from 4 mg IN in dogs (27) and 7.1-fold higher than reported from 4 mg IN dosed in humans (Table 1. and ref. 4).

**Fig 1.**
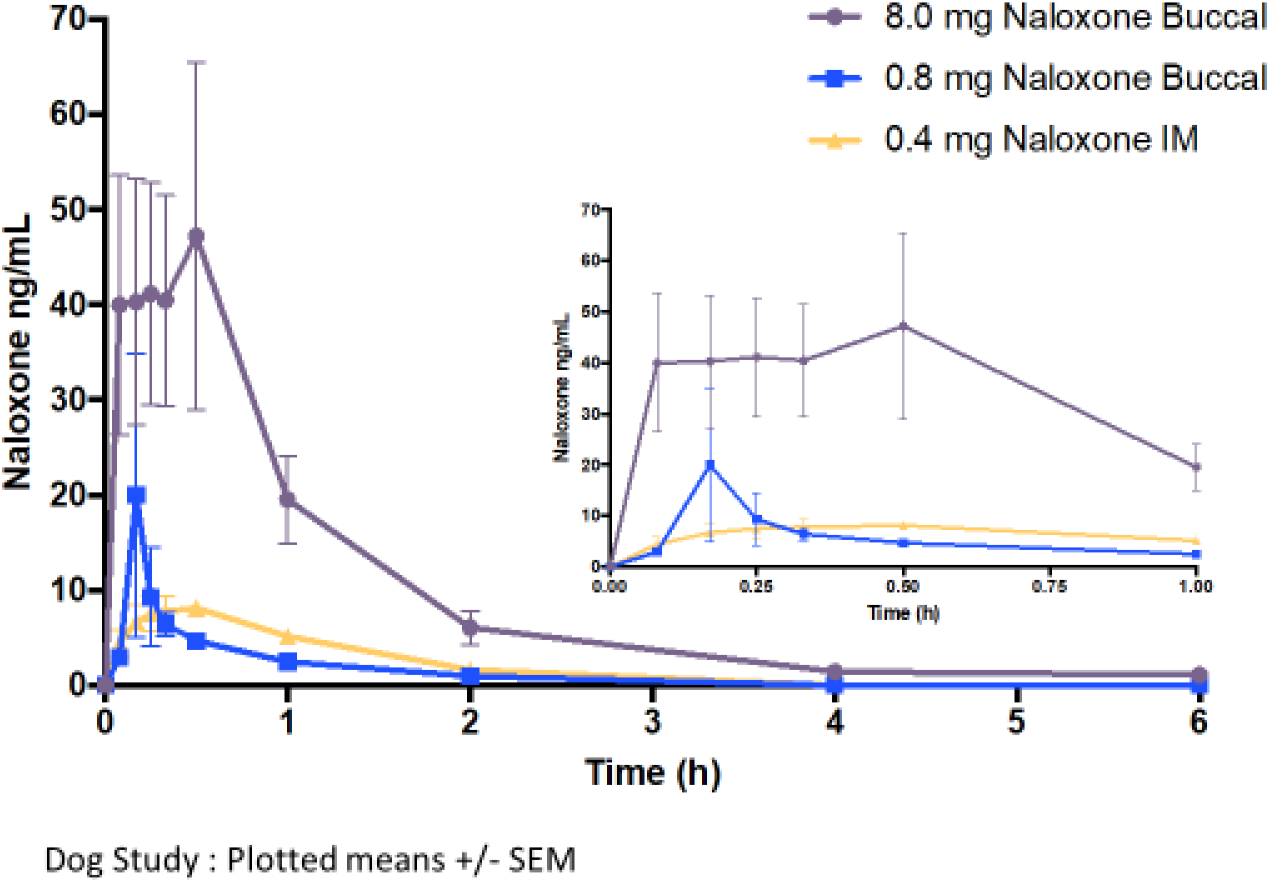
Mean +/− S.E.M plasma naloxone concentrations immediately before (baseline; 0 minutes) and at various time points following transbuccal swab administered naloxone (0.8 and 8.0 mg) and 0.4 mg IM injected to 3 healthy dogs in random crossover design involving 3 experimental periods (n=3 dogs/administration route/period) separated by 1 wk washout period. *Inset* displays mean plasma concentrations from 0 to 60 mins after administration.

**Fig 2.**
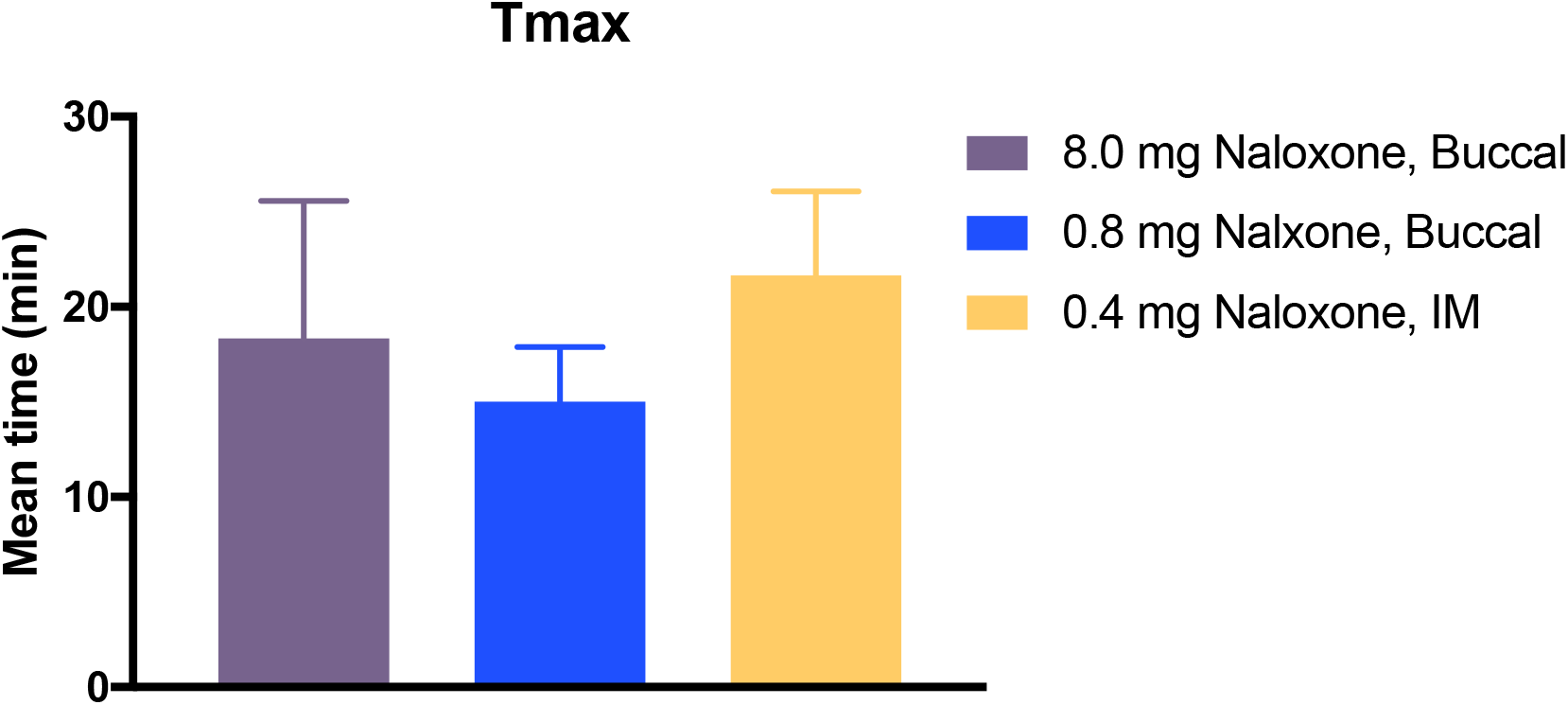
Mean time +/− S.E.M. to maximum plasma naloxone concentration after swab application of 0.8 and 8.0 mg on dog buccal mucosa compared with 0.4 mg IM injection.

**Fig 3.**
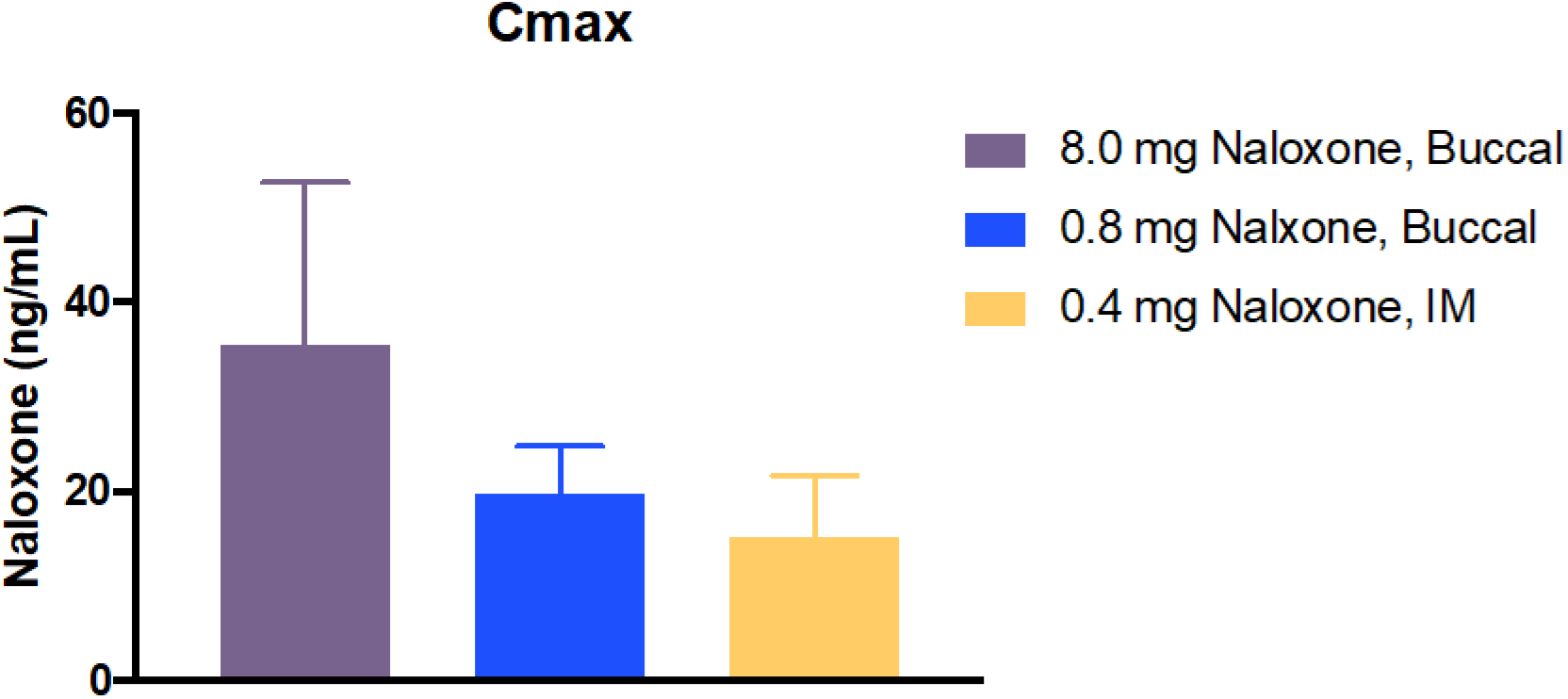
Mean +/− S.E.M of maximum plasma naloxone concentrations achieved after swab applications of 0.8 and 8.0 mg on dog buccal mucosa compared with 0.4 mg IM

**Fig 4.**
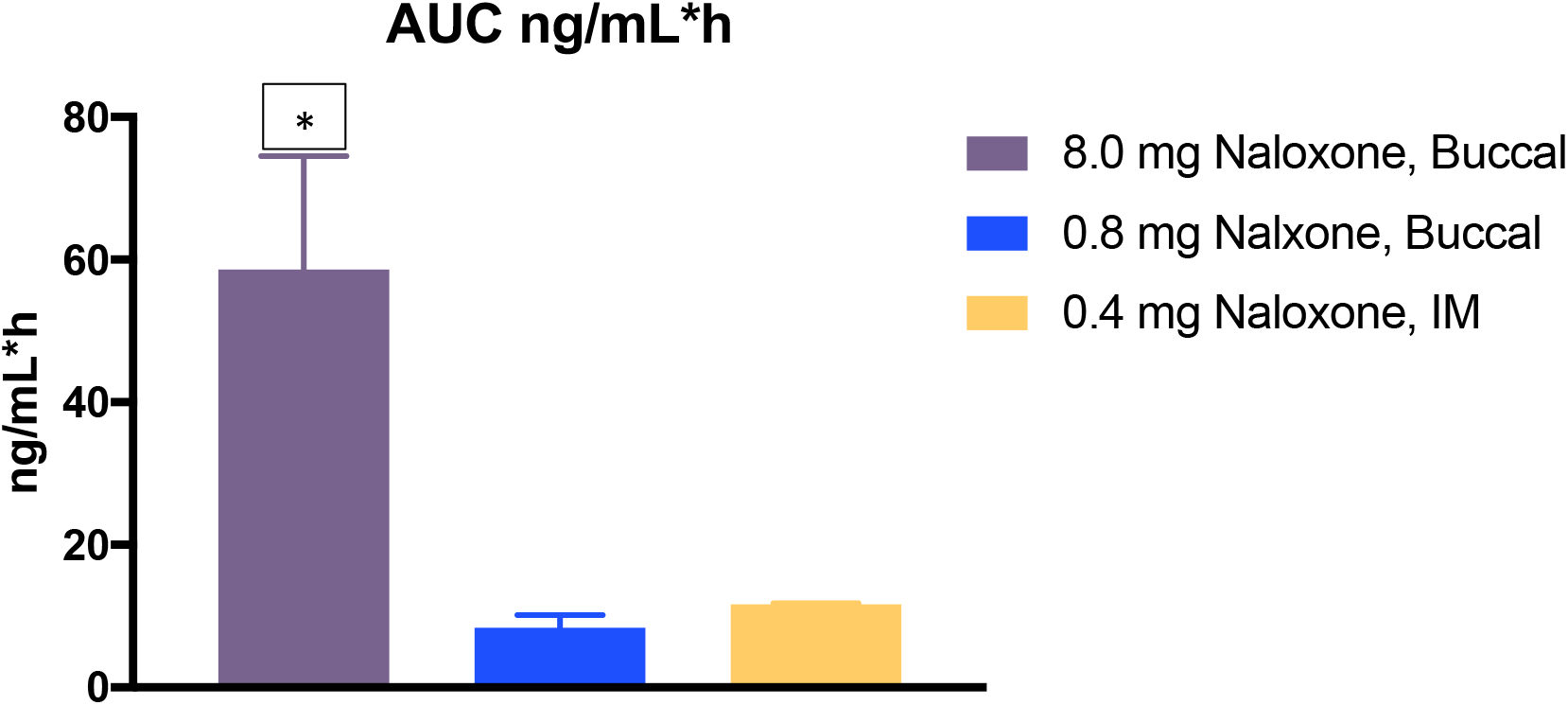
Mean +/− S.E.M of systemic exposure AUC plasma naloxone achieved after swab applications of 0.8 and 8.0 mg on dog buccal mucosa compared with 0.4 mg IM

**Table 1.**
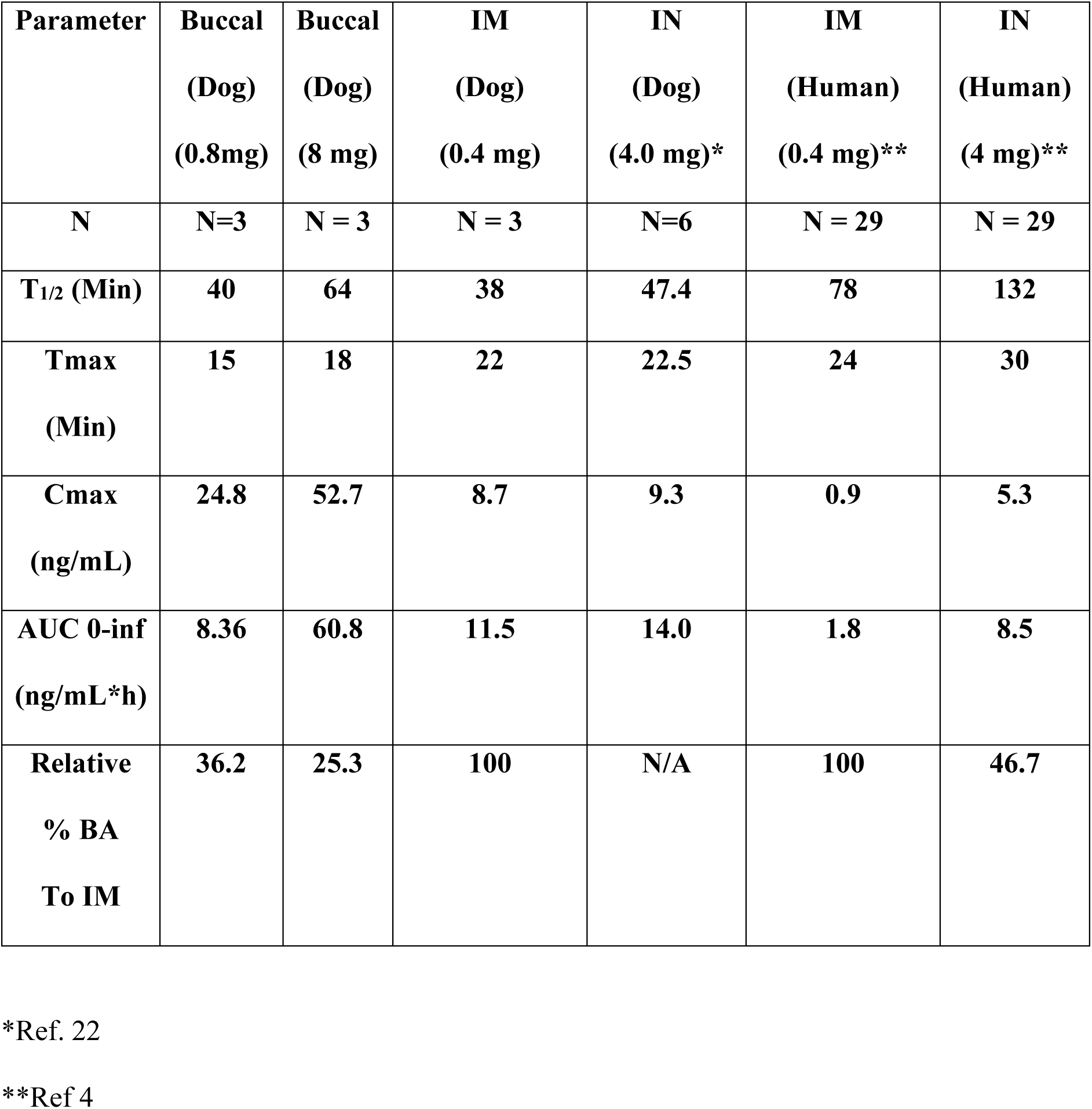
Comparison of PK Parameters for Naloxone Delivered by Buccal, IM and IN Routes of Administrations.

| **Parameter** | **Buccal**<br>**(Dog)**<br>**(0.8mg)** | **Buccal**<br>**(Dog)**<br>**(8 mg)** | **IM**<br>**(Dog)**<br>**(0.4 mg)** | **IN**<br>**(Dog)**<br>**(4.0 mg)\*** | **IM**<br>**(Human)**<br>**(0.4 mg)\*\*** | **IN**<br>**(Human)**<br>**(4 mg)\*\*** |
| --- | --- | --- | --- | --- | --- | --- |
| **N** | **N=3** | **N = 3** | **N = 3** | **N=6** | **N = 29** | **N = 29** |
| **T<sub>1/2</sub> (Min)** | **40** | **64** | **38** | **47.4** | **78** | **132** |
| **T<sub>max</sub>**<br>**(Min)** | **15** | **18** | **22** | **22.5** | **24** | **30** |
| **C<sub>max</sub>**<br>**(ng/mL)** | **24.8** | **52.7** | **8.7** | **9.3** | **0.9** | **5.3** |
| **AUC 0-inf**<br>**(ng/mL\*h)** | **8.36** | **60.8** | **11.5** | **14.0** | **1.8** | **8.5** |
| **Relative**<br>**% BA**<br>**To IM** | **36.2** | **25.3** | **100** | **N/A** | **100** | **46.7** |

\*Ref. 22
\*\*Ref 4

A comparison of PK summary data from dog buccal delivery of naloxone to referenced values (4) from dog IN, human IM and IN are shown in Table 1. Relative to IM bioavailability, buccal administration shows comparable blood levels to human IN administration. Similarly, rapid systemic absorption displayed by Tmax is comparable to human IM and IN naloxone administrations.

## Discussion

Deaths due to prescription opioids and illicit opioid overdose have been increasing at an alarming rate. Naloxone administration is the gold-standard antidote for emergency treatment of opioid overdoses. There are a variety of techniques and routes of administration used to deliver this life-saving drug including IM injections by syringes or autoinjectors and IN devices.^23–25^ Parenteral injections require technique and training that most individuals do not possess and even automatic injectors can be difficult to use possessing additional burdensome considerations such as accidental misfiring and needle sticks.^5–7^ For these and other reasons, emergency medical personnel in the field are reluctant to administer parenteral treatment. IN administration also possesses its own set of factors that can hinder intranasal drug absorption such as nasal congestion, lack of mucosa to absorb the drug due to surgery, bloody nose or other vasoconstrictors in drug use such cocaine abuse. These issues are further amplified with family members or caregivers using naloxone at home or laypersons applying treatment in public during emergencies that can affect timely administration. Even with approval of Narcan^®^ (nasal administration of naloxone HCl), many challenges remain even though FDA approval has allowed more widespread distribution and use. However there remains compelling demand for an easier-to-use, less cumbersome means of delivering naloxone with equivalent or superior performance characteristics.^6,24,25^

In this study, naloxone administered *via* the transbuccal route using a TOB gum resin formulation with attributes of film coating muco-adherence demonstrated maintenance contact with the mucous membrane of the buccal compartment to achieve rescue blood levels comparable to that obtained after IM injection of 0.4 mg naloxone, a benchmark efficacious dose *via* a parenteral route of administration. Certain botanical gums and polymers demonstrate interactions occurring through a variety of physical and chemical interactions in bonding to mucous membranes.^26–28^ Several drugs have been studied for orotransmucosal delivery by gum resins.^27,29^ The many advantages of buccal drug delivery, which is administration through the mucosal membranes lining the cheek (buccal mucosae), includes access to large surface area with high permeability close to a dense, shallow, vascular network facilitating rapid drug absorption conducive in the delivery of life-saving drugs such as naloxone. Furthermore, buccal administration is also be applicable to many other classes of drugs such as benzodiazepines used for anti-anxiety panic attacks, antinausea drugs such as scopolamine, prochlorperazine or ondansetron, anti-migraine triptans and perhaps even PDE-5 inhibitors for erectile dysfunction. It is even conceivable the delivery of peptides such as glucagon for emergency hypoglycemia or certain vaccine configurations could also be tested for advancement by transbuccal route of administration.^16,17^

This study demonstrates that a simple transbuccal swab application of naloxone in TOB has potential to provide additional and advantageous treatment route of administration for opioid overdosing among other therapeutic delivery applications. Understanding the widespread epidemic dangers of opioid usage and abuse it is conceivable that a buccal product could rapidly become the standard-of-care, deploying life-saving swabs in first aid kits, home medicine cabinets, personal cars, emergency vehicles, hospitals, schools, urgent care facilities, airplanes, restaurants or other public places where laypersons could simply administer the drug. Importantly, recent trends in rescuing opioid overdosing often times require multiple doses of naloxone to reach adequate blood levels due in part by the prevalence of more powerful and higher affinity synthetic opioids such as fentanyl.^7^ This simple buccal swabbing of naloxone demonstrating a rapid and sustained component with longer exposure coverage than IM and IN may also provide additional advantages to rescuing in this current surge of opioid overdosing worldwide.

## Acknowledgements

The authors thank Dr. Jan Scicinski for assistance in formatting, statistical analysis and proofing this manuscript.

## Notes

### Competing Interest Statement

The authors have declared no competing interest.

### Summary of Updates

Addition of references and subsequent reference citations throughout manuscript.

